# Integrated Evaluation of Osmotic and Antioxidant Defense Mechanisms in Cotton Genotypes Exposed to NaCl Stress

**DOI:** 10.64898/2026.06.03.729956

**Authors:** Nodira R. Rakhmatova, Azadakhan S. Imamkhodjaeva, Ilxom B. Salahutdinov, Venera S. Kamburova, Shakhnoza B. Kadirova, Farxod S. Radjapov, Jurabek K. Norbekov, Muyassar Zakirova, Zebo Z. Yuldashova, Rasul A. Jumaev, Zabardast T. Buriev

## Abstract

Salinity stress is one of the major abiotic factors limiting cotton productivity worldwide by inducing osmotic imbalance, oxidative stress, and metabolic disturbances in plant tissues. The present study aimed to evaluate the physiological and biochemical responses of different cotton (*Gossypium hirsutum* L.) genotypes under NaCl-induced salinity stress through analysis of proline accumulation, antioxidant enzyme activities, and lipid peroxidation intensity.

The experiment was conducted under controlled conditions using several cotton genotypes exposed to different NaCl concentrations. Proline content, superoxide dismutase (SOD), catalase (CAT), and malondialdehyde (MDA) levels were analyzed as major biochemical indicators associated with salinity tolerance and oxidative stress responses. In addition, modern bubble heatmap visualization was applied for comparative assessment of genotype-specific stress response patterns under saline treatments.

The obtained results demonstrated that increasing NaCl concentrations generally stimulated proline accumulation and enhanced antioxidant enzyme activities in most investigated cotton genotypes. Increased SOD and CAT activities indicated activation of enzymatic antioxidant defense mechanisms under salinity stress conditions. Simultaneously, elevated MDA accumulation reflected enhanced oxidative membrane damage caused by excessive reactive oxygen species (ROS) production under saline environments.

Considerable genotype-dependent variability was observed among the investigated cotton varieties. Genotypes such as “Nasaf”, “Gulbahor-2”, “Ravnaq-1”, “Buxoro-6”, “Afsona”, “Baraka”, “Namangan-77”, “Porloq-1”, and “C-4727” demonstrated comparatively stronger physiological and antioxidant responses under salinity stress conditions, suggesting relatively higher adaptive capacity to NaCl-induced stress.

The heatmap visualization additionally confirmed substantial heterogeneity among cotton genotypes in biochemical stress responses and allowed comprehensive comparative interpretation of salinity-induced physiological variability.

Overall, the present findings suggest that proline accumulation, antioxidant enzyme activities (SOD and CAT), and MDA content may serve as important biochemical markers for evaluation of salinity tolerance in cotton. The identified stress-tolerant genotypes may therefore represent valuable genetic resources for future breeding programs aimed at improving cotton productivity under saline environmental conditions.

## Introduction

Salinity stress is considered one of the most serious abiotic factors limiting agricultural productivity worldwide. Increasing soil salinization negatively affects plant growth, development, physiological processes, and crop yield, particularly in arid and semi-arid regions where irrigation practices and climate change accelerate salt accumulation in agricultural soils. According to recent estimates, more than 20% of irrigated agricultural lands worldwide are affected by salinity stress, and this problem continues to expand annually (Hasanuzzaman et al., 2021; Wang et al., 2024).

Cotton (*Gossypium hirsutum* L.) is one of the most economically important fiber crops globally and plays a major role in the textile industry and agricultural economy of many countries. Although cotton is considered relatively more tolerant to salinity compared with several other field crops, excessive salt accumulation significantly reduces seed germination, plant growth, photosynthetic efficiency, and fiber productivity (Maryum et al., 2022; Ullah et al., 2026).

Salinity stress induces osmotic imbalance, ionic toxicity, nutrient disorders, and excessive production of reactive oxygen species (ROS) in plant cells. Overaccumulation of ROS such as superoxide radicals, hydrogen peroxide, and hydroxyl radicals may cause oxidative damage to cellular membranes, proteins, pigments, enzymes, and nucleic acids (Foyer and Noctor, 2005; Tripathy and Oelmüller, 2012). Therefore, regulation of oxidative stress represents one of the major physiological mechanisms associated with plant adaptation to saline environments.

Plants have evolved complex defense systems to protect cellular structures against ROS-induced oxidative damage. These defense mechanisms include enzymatic antioxidant systems such as superoxide dismutase (SOD), catalase (CAT), and peroxidases, as well as non-enzymatic antioxidant compounds and compatible osmolytes (Gill and Tuteja, 2010; Hasanuzzaman et al., 2020).

Superoxide dismutase is considered the first enzymatic barrier against oxidative stress because it catalyzes the conversion of superoxide radicals into hydrogen peroxide and molecular oxygen. Catalase subsequently detoxifies hydrogen peroxide into water and oxygen, thereby reducing oxidative membrane injury under stress conditions (Noreen and Ashraf, 2009; Hasanuzzaman et al., 2021).

Accumulation of compatible osmolytes such as proline also represents an important adaptive mechanism under salinity stress. Proline contributes to osmotic adjustment, stabilization of proteins and membranes, maintenance of cellular water balance, and protection against oxidative damage (Ashraf, 2009; Hosseinifard et al., 2022). Enhanced proline biosynthesis under saline conditions has been widely reported in cotton and other crop species exposed to abiotic stress (Meloni et al., 2001; Zulfiqar and Ashraf, 2023).

Malondialdehyde (MDA), a major product of membrane lipid peroxidation, is widely used as a biochemical indicator of oxidative stress intensity and cellular membrane damage in plants subjected to salinity stress (Czégény et al., 2014; Wang et al., 2024). Increased MDA accumulation generally reflects enhanced oxidative injury caused by excessive ROS production under stress conditions.

Previous studies have demonstrated that salinity tolerance in cotton is closely associated with efficient osmotic adjustment and antioxidant defense regulation (Sekmen et al., 2014; Ahmad et al., 2010). However, physiological and biochemical responses to salinity stress may vary considerably among cotton genotypes depending on their genetic background and stress adaptation capacity.

Therefore, identification of salt-tolerant cotton genotypes based on biochemical stress markers represents an important objective for development of stress-resilient cultivars suitable for saline agricultural regions.

The present study aimed to evaluate the physiological and biochemical responses of different cotton genotypes under NaCl-induced salinity stress by analyzing proline accumulation, antioxidant enzyme activities (SOD and CAT), and MDA content. In addition, modern bubble heatmap visualization was applied to comparatively assess genotype-specific stress response patterns under different salinity treatments.

According to Babadjanova, et al. (2025), abiotic and biotic stresses are major global challenges that negatively affect plant growth, development, physiological processes, agricultural productivity, and global food security by disrupting photosynthesis, water balance, and antioxidant defense systems.

## Materials and Methods

The plant material consisted of 28 cotton cultivars: Afsona, Sulton, Namangan-34, Baraka, Kupaysin, Gulbahor-2, C-4727, Namangan-102, Turkan, Namangan-76, Nasaf, Xin Lu Zao-78, Porloq-1, Omad, Chuntay-2, Ravnaq-1, Jinken-1402, Ishonch, TM-1, Buxoro-102, Xin Lu Zhong-87, Chimboy, Junjen, Buxoro-10, Buxoro-6, Buxoro-14, Navbahor-2 and Kelejak.

The experiment was conducted under controlled phytotron conditions. To simulate salt stress, plants were divided a control group and four salt-treated groups (50, 100, 150, and 200 mM NaCl). Both control and salt-treated plants were irrigated for 21 days. Control plants received water (0 mM NaCl), whereas salt-treated plants were received NaCl solutions at the respective concentrations.

After the treatment period, fully expanded leaves were harvested for biochemical analyses. The content of malondialdehyde (MDA) in fresh leaf tissue, as well as the activities of superoxide dismutase (SOD) and catalase (CAT), were determined spectrophotometrically and expressed as nmol g⁻¹ fresh weight.

Each genotype was grown under five NaCl treatments (0, 50, 100, 150, and 200 mM), with five biological replicates per treatment (Rakhmatova et al., 2025).

### Determination of malondialdehyde (MDA) content

Malondialdehyde (MDA) content in plant samples was determined according to a modified method of Heath and Packer (1968). Briefly, 100 mg of fresh plant material was homogenized in 2 mL of 20% (w/v) trichloroacetic acid (TCA). The homogenate was centrifuged at 10,000 × g for 15 min at 4 °C.

After centrifugation, 0.5 mL of the supernatant was mixed with 1.5 mL of 0.5% (w/v) thiobarbituric acid (TBA) prepared in 20% TCA. The reaction mixtures were incubated in a water bath at 95 °C for 30 min and then rapidly cooled in an ice bath. To correct for nonspecific absorbance caused by sample pigmentation, a control sample without TBA was prepared by adding 1.5 mL of 20% TCA to 0.5 mL of the supernatant.

Absorbance of both control and TBA-treated samples was measured at 532 nm and 600 nm using a Synergy™ NT microplate reader (Bio Tek Instruments, USA).

MDA concentration was calculated using the following equation:

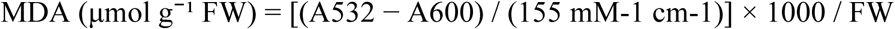

where: A532 and A600 are absorbance values at 532 and 600 nm, respectively; 155 mM-1 cm-1 is the extinction coefficient of the TBA–MDA complex; and FW is the fresh weight of the sample (g).

### Superoxide dismutase (SOD) determination

The activity of the enzyme SOD was determined based on the inhibition of superoxide radical formation during the autoxidation of adrenaline in an alkaline medium in vitro at a wavelength of 347 nm with some modifications (Leonowicz et al., 2018). Briefly, 0.1 mL of distilled water and 0.1 mL of 0.1 % (5.46 mM) epinephrine hydrochloride solution were added to 2 mL of 0.2 M bicarbonate buffer (pH = 10.65), thoroughly and rapidly mixed and placed in a Cary UV-60 spectrophotometer. Optical density was recorded every 30 sec for 5 min at 347 nm using a 10 mm quartz cuvette. After that, 0.1 mL of the enzyme source (vegetable homogenate) and 0.1 mL of 0.1 % epinephrine hydrochloride were added to 2 mL of buffer (pH = 10.65), mixed and the optical density was measured. The homogenate for determining SOD activity was prepared as follows: 100 mg of plant leaves were ground in a porcelain mortar with 1 mL of 10 mM Tris-HCl buffer (pH 7.8). The homogenate was centrifuged at 7000 g for 15 min at 4 °C and the resulting supernatant was used as the enzyme source.

SOD activity is calculated using the following formula:

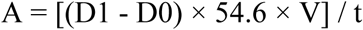

Where, A is the activity of SOD, mM of decomposed adrenaline per min, D1 is the optical density of the experimental sample, D0 is the optical density of the control sample, 54.6 mM is the concentration of adrenaline in the cuvette, V is the source of the enzyme in the sample, t is the inhibition time of autoxidation of adrenaline (5 min).

### Catalase (CAT) determination

Catalase (CAT) activity was measured following the method of Beers and Sizer (1952), with minor modifications. The reaction mixture (1.5 mL) consisted of 100 mM phosphate buffer (pH 7.0), 0.1 mM EDTA, 20 mM H2O2, and 50 µL of enzyme extract. The reaction was initiated by adding the enzyme extract, and the decrease in H₂O₂ concentration was monitored spectrophotometrically at 240 nm using a molar extinction coefficient of 36 M-1 cm-1. Enzyme activities were expressed as units per milligram of protein (EU mg-1 protein).

## Statistical analysis

### Materials and Methods: Analysis of Physiological and Biochemical Parameters and Statistical Visualization

Proline accumulation, malondialdehyde (MDA) content, as well as the activities of the antioxidant enzymes superoxide dismutase (SOD) and catalase (CAT) in cotton genotypes under NaCl-induced salt stress were evaluated using spectrophotometric analytical methods followed by statistical processing and graphical visualization of the obtained data. Experimental data obtained under different NaCl concentrations (control, 50, 100, 150, and 200 mM NaCl) were analyzed to identify genotype-specific physiological and biochemical responses of cotton plants to salinity stress.

For each genotype and treatment variant, mean values and standard errors were calculated based on repeated experimental measurements. Statistical data processing was performed using Microsoft Excel and Python-based analytical tools. The obtained numerical datasets were organized into matrices reflecting changes in proline content, MDA level, and antioxidant enzyme activities depending on the severity of salt stress.

For scientific visualization and comparative analysis, the Python programming environment with the Pandas and Matplotlib libraries was used. The Pandas library was applied for data processing, structuring, and statistical organization, whereas Matplotlib was used to generate publication-quality graphs and visualizations.

To achieve a more detailed interpretation of the physiological and biochemical responses of cotton genotypes, several modern data visualization approaches were employed. The selective multi-line visualization approach was used to display the most contrasting genotype responses to salt stress. Within this approach, genotypes characterized by high antioxidant enzyme activity, increased proline accumulation, and relatively low levels of lipid peroxidation (MDA) were highlighted, thereby improving graph readability and emphasizing the most pronounced physiological and biochemical changes.

Average response curve analysis was performed to evaluate the overall response dynamics of cotton genotypes to increasing NaCl concentrations. For each salinity level, average values of proline content, MDA level, and SOD and CAT activities were calculated across all investigated genotypes, allowing visualization of the general trends in antioxidant and osmoprotective responses under salt stress conditions.

Statistical analyses were performed using one-way ANOVA followed by Tukey’s multiple comparison test at p < 0.05. The results are presented as mean ± standard error (SE).

To provide a concise overview of the experimental design, investigated cotton genotypes, salinity treatments, biochemical parameters, and statistical approaches applied in the present study, the main methodological framework is summarized in Table 1.

**Table 1.** Experimental design, investigated parameters, and statistical approaches used in the study.

| Experimental factors | Description |
| --- | --- |
| Plant material | Cotton (*Gossypium hirsutum* L.) genotypes |
| Investigated genotypes | Nasaf, Gulbahor-2, Ravnaq-1, Buxoro-6, Afsona, Baraka, Namangan-77, Porloq-1, C-4727 and other studied genotypes |
| Stress treatment | NaCl-induced salinity stress |
| Salinity levels | Control, 100 mM NaCl, 150 mM NaCl, 200 mM NaCl |
| Experimental conditions | Controlled laboratory/growth chamber conditions |
| Experimental design | Completely randomized design |
| Number of replications | Three biological replicates |
| Investigated parameters | Proline, SOD, CAT, and MDA |
| Statistical analysis | Mean ± standard deviation (SD), ANOVA, significance at $p < 0.05$ |
| Visualization approach | Integrated heatmap visualization and statistical analysis of physiological and biochemical responses |

## 1. RESULTS

### 1.1. Effect of NaCl stress on proline accumulation in cotton genotypes

The obtained results demonstrated substantial genotypic variability in proline accumulation with increasing NaCl concentration, indicating differences in the physiological responses of cotton genotypes to salt stress conditions. Proline is widely regarded as an important osmoprotective metabolite involved in osmotic regulation, stabilization of cellular membranes, protection of proteins, and neutralization of reactive oxygen species under abiotic stress conditions. Consequently, the observed increase in proline content under salt treatment was accompanied by the activation of protective responses in the investigated genotypes.

Under control conditions, most genotypes were characterized by relatively low basal levels of proline, indicating the absence of pronounced osmotic stress and the normal progression of physiological and biochemical processes. However, increasing NaCl concentrations (50–200 mM) resulted in a significant enhancement of proline accumulation in several genotypes, although the degree of response differed considerably among cultivars.

Particularly high proline accumulation was observed in the genotypes “Gulbahor-2”, “Nasaf”, “Buxoro-6”, “Ravnaq-1”, “Afsona”, “Baraka”, “Namangan-77”, “Porloq-1”, and “C-4727”. The genotype “Gulbahor-2” exhibited a sharp increase in proline content already at 50 mM NaCl, reaching one of the highest values among all investigated treatments. In the genotype “Nasaf”, maximum proline accumulation was observed under 100–150 mM NaCl treatment.

The genotypes “Buxoro-6” and “Ravnaq-1” were characterized by a gradual and stable increase in proline content with increasing salt concentration, reaching maximum values at 200 mM NaCl. A similar tendency was observed in the genotypes “Afsona”, “Baraka”, “Namangan-77”, and “Porloq-1”, which also demonstrated high levels of proline accumulation under moderate and severe salinity conditions. The genotype “C-4727” was characterized by a stable enhancement of proline biosynthesis with increasing NaCl concentration.

In contrast, the genotypes “Omad”, “Xin Lu Zhou-78”, and “Chuntay-2” demonstrated comparatively low proline accumulation under salt stress conditions.

The heatmap (Fig. 1) also confirmed the pronounced heterogeneity in the responses of different genotypes to salt stress. A clear distribution of cultivars with high and low levels of proline accumulation reflected differences in the intensity of responses to NaCl exposure.

**Figure 1.**
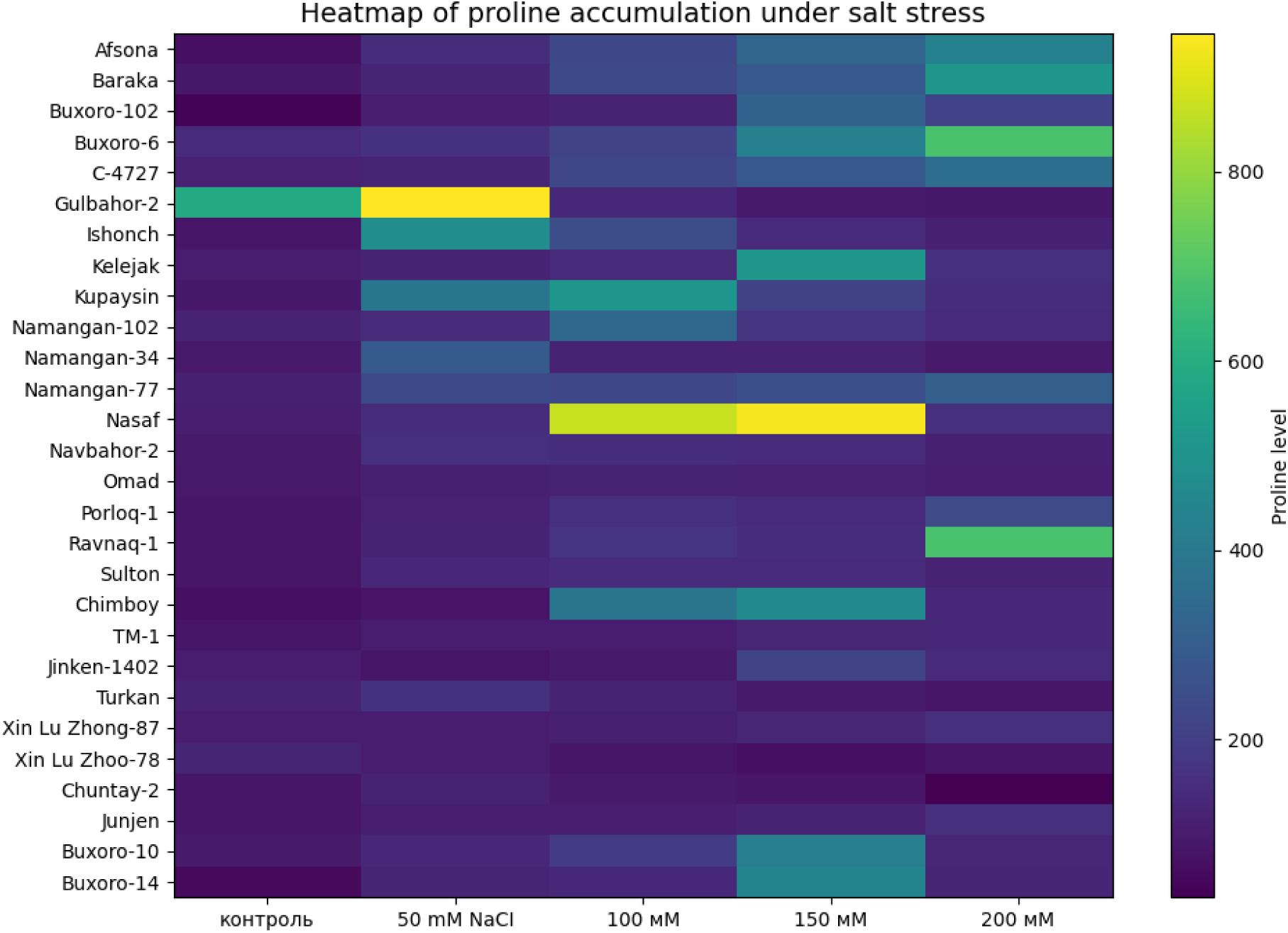
Heatmap of proline accumulation in cotton genotypes under salt stress conditions.

Genotypes characterized by more intense coloration under high NaCl concentrations corresponded to elevated levels of proline accumulation.

Overall, the results of the study demonstrated that salt stress significantly stimulated proline biosynthesis in the majority of the investigated cotton genotypes; however, the extent of accumulation depended on both genotype and salinity level.

### 1.2. Effect of NaCl stress on catalase activity in cotton genotypes

Figure 2 presents a heatmap visualization of catalase (CAT) activity in different cotton genotypes under NaCl-induced salt stress conditions. The analysis demonstrated that increasing NaCl concentrations stimulated the activation of antioxidant defense mechanisms in the majority of the studied genotypes. A particularly pronounced increase in CAT activity was observed under 150 and 200 mM NaCl treatments, indicating enhanced detoxification of reactive oxygen species (ROS) generated during oxidative stress.

**Figure 2.**
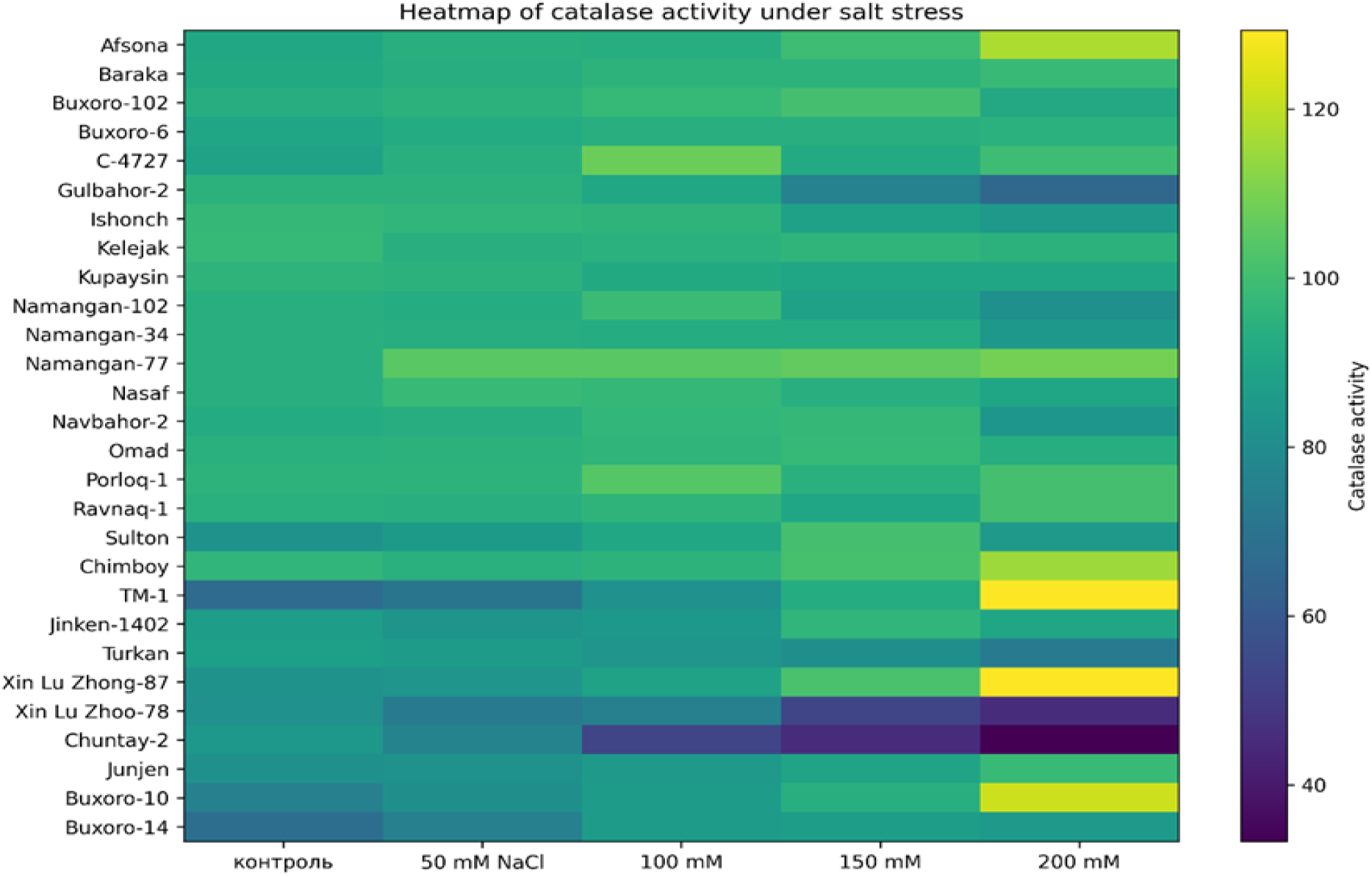
Heatmap visualization of catalase (CAT) activity in cotton genotypes under salt stress conditions.

Heatmap analysis revealed that several genotypes exhibited remarkably high antioxidant activity under saline conditions. In particular, the TM-1 genotype showed the highest catalase activity at 200 mM NaCl. Similarly, significant increases in CAT activity under severe salt stress were observed in Xin Lu Zhong-87, Buxoro-10, Afsona, Baraka, and Namangan-77 genotypes. These findings suggest that these genotypes possess a stronger ability to protect cellular structures against oxidative damage under stress conditions.

Furthermore, the Nasaf and Kupaysin genotypes maintained relatively high CAT activity under 100–150 mM NaCl treatments, indicating a stable antioxidant defense system and efficient physiological adaptation to salinity stress.

In contrast, the Chuntay-2 and Xin Lu Zhoo-78 genotypes exhibited reduced CAT activity at elevated NaCl concentrations. The substantial decline in enzyme activity under 200 mM NaCl suggests a weaker antioxidant defense response and greater sensitivity to salt stress.

Additionally, the Ravnaq-1, C-4727, and Buxoro-6 genotypes maintained comparatively high CAT activity even under severe salinity conditions, indicating their potential as salt-tolerant genotypes (Figure 2).

Overall, the obtained results confirm the crucial role of catalase as one of the key antioxidant enzymes involved in cotton adaptation to saline environments. The heatmap visualization clearly demonstrated physiological and biochemical differences among the studied genotypes and identified promising candidates for future breeding programs aimed at improving salt tolerance in cotton.

### 1.3. Effect of NaCl stress on superoxide dismutase activity in cotton genotypes

The figure 3 presents a heatmap visualization of superoxide dismutase (SOD) activity in different cotton genotypes under NaCl-induced salt stress conditions. The obtained results demonstrated that salinity stress significantly affected SOD activity, while the response pattern varied depending on genotype and NaCl concentration.

**Figure 3.**
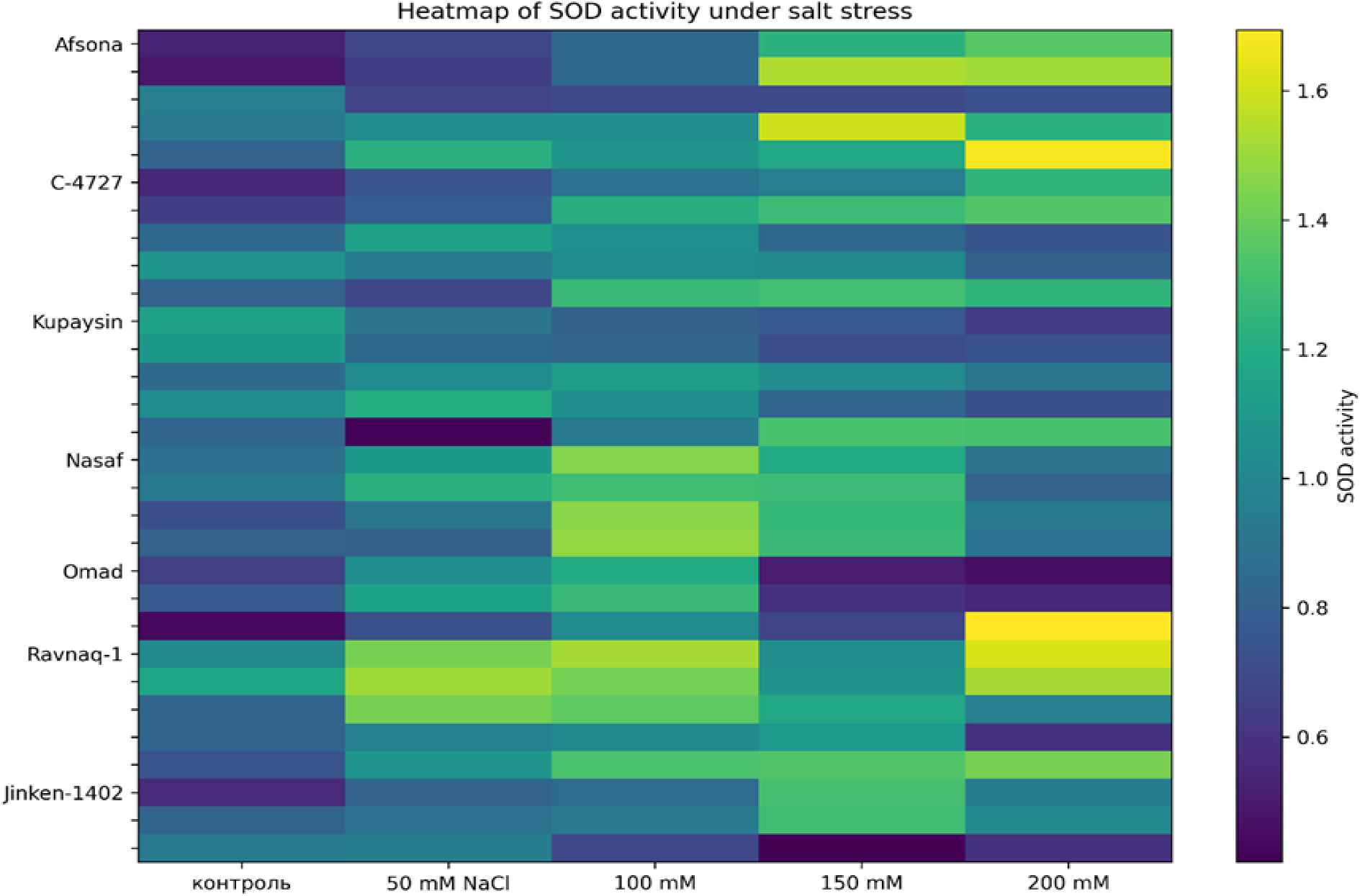
Heatmap visualization of superoxide dismutase (SOD) activity in cotton genotypes under salt stress conditions.

Under control conditions, most genotypes exhibited relatively low to moderate SOD activity. However, increasing NaCl concentrations generally stimulated SOD activity in the majority of the studied genotypes. A particularly pronounced increase in enzyme activity was observed under 100–200 mM NaCl treatments, indicating activation of antioxidant defense mechanisms against reactive oxygen species (ROS) generated under oxidative stress conditions.

Heatmap analysis revealed that several genotypes exhibited remarkably high SOD activity. In particular, the genotype “Ravnaq-1” showed the highest SOD activity at 100 and 200 mM NaCl treatments, suggesting strong antioxidant defense capacity under salinity stress.

Similarly, the “Nasaf” genotype demonstrated a significant increase in SOD activity under 100–150 mM NaCl conditions. In the “Afsona” genotype, SOD activity gradually increased with increasing salinity level, reaching high values under 150–200 mM NaCl treatments. This finding may indicate effective physiological adaptation of this genotype to saline environments.

Furthermore, the genotypes “Jinken-1402”, “C-4727”, and “Kupaysin” also exhibited moderate to high SOD activity under salt stress conditions. Particularly, “C-4727” maintained relatively stable enzyme activity across different NaCl concentrations, suggesting physiological stability and adaptive tolerance to salinity stress.

In contrast, the genotype “Omad” exhibited decreased SOD activity under high salinity conditions. Especially under 150–200 mM NaCl treatments, the reduced enzyme activity may indicate weaker antioxidant defense mechanisms and greater sensitivity to salt stress. In some genotypes, a temporary decline in SOD activity was observed at 50 mM NaCl, followed by reactivation at higher salinity levels.

The heatmap visualization clearly demonstrated substantial genotype-dependent variability in SOD activity. Increased color intensity corresponded to higher enzyme activity and reflected more efficient antioxidant defense responses in salt-tolerant genotypes.

Overall, the obtained results confirm the important role of superoxide dismutase (SOD) in protecting cotton plants against salt-induced oxidative stress. Genotypes characterized by elevated SOD activity may serve as valuable genetic resources for future breeding programs aimed at developing salt-tolerant cotton varieties.

### 1.4. Effect of NaCl stress on malondialdehyde (MDA) content in cotton genotypes

The figure 4 presents a heatmap visualization of malondialdehyde (MDA) accumulation in different cotton genotypes under NaCl-induced salt stress conditions. MDA is one of the major products of lipid peroxidation, and its accumulation reflects oxidative damage to cellular membranes. The obtained results demonstrated that salinity stress significantly affected MDA accumulation, while the response pattern varied depending on genotype and NaCl concentration.

**Figure 4.**
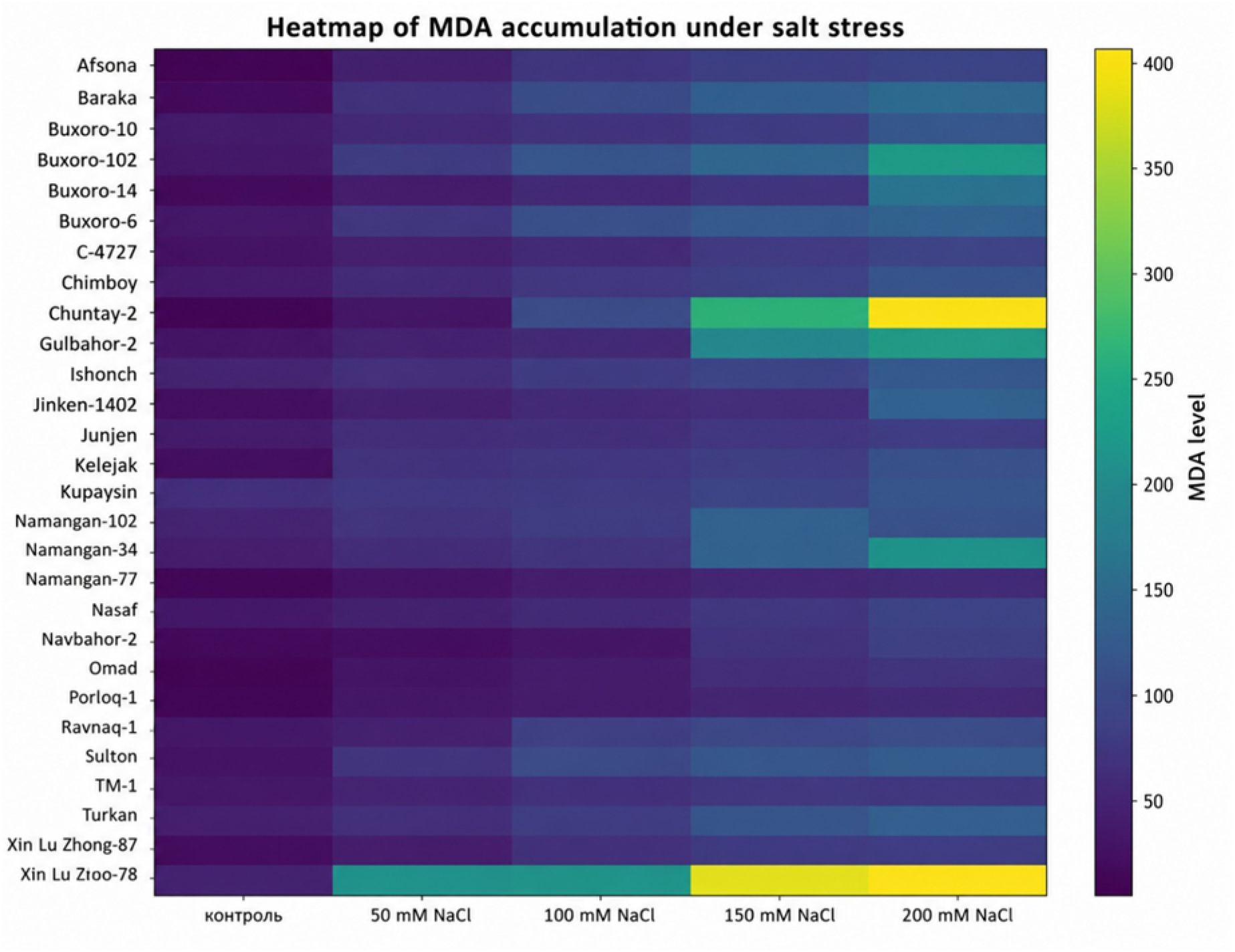
Heatmap visualization of malondialdehyde (MDA) accumulation in cotton genotypes under salt stress conditions.

Under control conditions, most genotypes exhibited relatively low MDA content. However, gradual increases in NaCl concentration led to elevated MDA levels in the majority of the studied genotypes. A particularly sharp increase in MDA accumulation was observed under 150–200 mM NaCl treatments, indicating severe oxidative stress and membrane damage under high salinity conditions.

Heatmap analysis revealed that the genotype “Xin Lu Zhoo-78” exhibited the highest MDA accumulation among all studied genotypes. In this genotype, MDA levels increased markedly beginning from 50 mM NaCl and reached maximum values under 150–200 mM NaCl treatments. This finding suggests high sensitivity of this genotype to salt stress and insufficient efficiency of antioxidant defense mechanisms.

Similarly, the genotype “Chuntay-2” also showed substantial increases in MDA accumulation under 100–200 mM NaCl conditions. Particularly high MDA content under 200 mM NaCl indicates intensive lipid peroxidation and severe membrane damage. The genotypes “Gulbahor-2”, “Namangan-34”, and “Buxoro-102” also demonstrated significant increases in MDA levels under elevated salinity conditions.

Moderate increases in MDA accumulation were observed in the genotypes “Turkan”, “Ravnaq-1”, “Sulton”, “Chimboy”, and “Jinken-1402”, suggesting intermediate physiological tolerance to salt stress. In particular, “Ravnaq-1” and “Sulton” maintained relatively stable MDA levels, indicating partial adaptation to saline environments.

In contrast, the genotypes “Afsona”, “Nasaf”, “Navbahor-2”, “Omad”, “Porloq-1”, “Buxoro-10”, and “Buxoro-14” maintained comparatively low MDA levels even under high NaCl concentrations. This suggests enhanced membrane stability and efficient antioxidant defense mechanisms in these genotypes. Particularly low MDA accumulation was observed in “Afsona” and “Nasaf”, indicating superior salt tolerance.

The heatmap visualization clearly demonstrated substantial genotype-dependent variability in MDA accumulation patterns. Increased color intensity corresponded to higher MDA levels and reflected enhanced oxidative damage under salinity stress conditions.

Overall, the obtained results confirm that NaCl stress intensifies lipid peroxidation processes in cotton genotypes. Lower MDA accumulation appears to be closely associated with effective antioxidant defense systems and enhanced salt tolerance. Therefore, genotypes such as “Afsona”, “Nasaf”, “Porloq-1”, and “Buxoro-14” may serve as promising genetic resources for breeding salt-tolerant cotton varieties.

Figure 5 presents an integrated heatmap analysis and comparative evaluation of biochemical responses in cotton genotypes under NaCl-induced salinity stress. The visualization summarizes the changes in antioxidant enzyme activities (SOD and CAT), proline accumulation, and MDA content across different salinity levels, allowing rapid identification of genotype-dependent stress tolerance patterns. In addition, the integrated tolerance index at 200 mM NaCl provides a comparative assessment of salt tolerance among the studied genotypes based on combined biochemical indicators.

**Figure 5.**
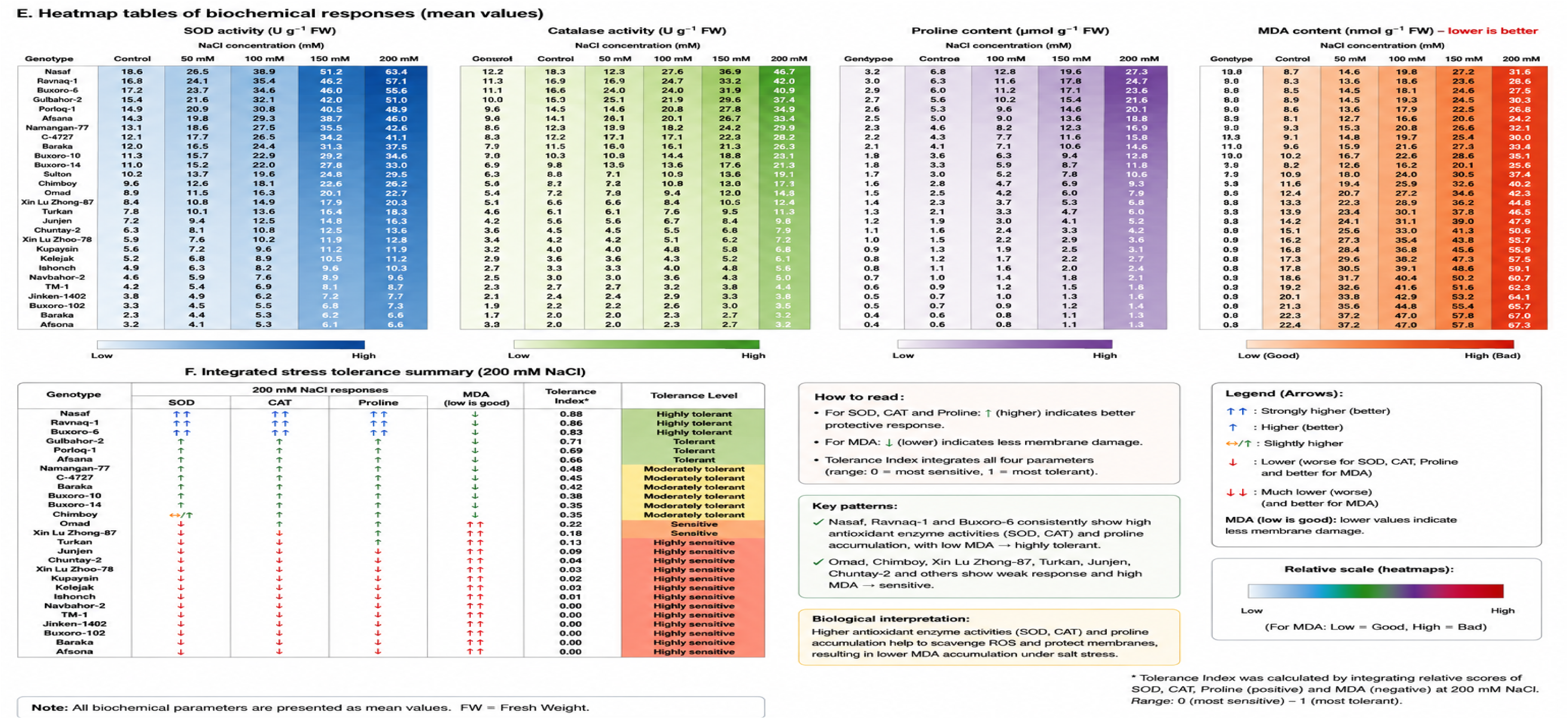
Integrated heatmap visualization of physiological and biochemical responses of cotton genotypes under NaCl-induced salinity stress. The figure summarizes genotype-dependent variations in superoxide dismutase (SOD) activity, catalase (CAT) activity, proline accumulation, and malondialdehyde (MDA) content under different NaCl concentrations. Increased SOD and CAT activities together with enhanced proline accumulation and relatively lower MDA content indicate improved salinity tolerance and more efficient antioxidant defense mechanisms.

The integrated heatmap visualization (Figure 5) demonstrated clear genotype-dependent variability in the physiological and biochemical responses of cotton genotypes under NaCl-induced salinity stress. In general, increasing salinity levels stimulated antioxidant defense activity and osmoprotective responses; however, the intensity and coordination of these reactions differed considerably among genotypes.

Salt-tolerant genotypes such as “Nasaf”, “Gulbahor-2”, “Ravnaq-1”, “Buxoro-6”, “Porloq-1”, and “Afsona”, as well as relatively tolerant genotypes including “Baraka” and “Namangan-77”, were characterized by coordinated increases in SOD and CAT activities together with enhanced proline accumulation under moderate and severe salinity conditions. Although MDA content also increased under stress conditions, its accumulation remained comparatively lower and more controlled than in sensitive genotypes. This pattern suggests more efficient ROS detoxification, improved membrane stability, and stronger adaptive capacity under saline environments.

In contrast, comparatively salt-sensitive genotypes such as “Xin Lu Zhoo-78”, “Chuntay-2”, and “Omad” exhibited weaker antioxidant responses accompanied by relatively higher MDA accumulation under elevated NaCl concentrations. Excessive MDA accumulation in these genotypes may indicate enhanced membrane lipid peroxidation and insufficient antioxidant protection against oxidative stress.

The integrated visualization further demonstrated that increased MDA accumulation under salinity stress does not necessarily contradict stress tolerance. Moderate elevation of MDA was observed even in tolerant genotypes because salinity-induced oxidative stress inevitably promotes ROS formation. However, tolerant genotypes maintained a more balanced relationship between antioxidant enzyme activity and oxidative damage, whereas sensitive genotypes exhibited disproportionately high MDA accumulation combined with weaker antioxidant defense responses. Overall, the combined analysis of SOD, CAT, proline, and MDA responses indicates that efficient antioxidant regulation together with osmotic adjustment plays a critical role in salinity tolerance in cotton genotypes. Therefore, the identified tolerant genotypes may serve as valuable genetic resources for future breeding programs aimed at improving cotton adaptation to saline environments.

Figure 6 illustrates the average physiological and biochemical responses of cotton genotypes under increasing NaCl concentrations. The integrated visualization demonstrates dynamic changes in proline accumulation, antioxidant enzyme activities (SOD and CAT), and MDA content, highlighting the coordinated activation of osmoprotective and antioxidant defense mechanisms under salinity stress. In general, most protective parameters reached their maximum levels at 150 mM NaCl, whereas MDA accumulation continued to increase at higher salinity levels, indicating enhanced membrane damage under severe stress conditions.

**Figure 6.**
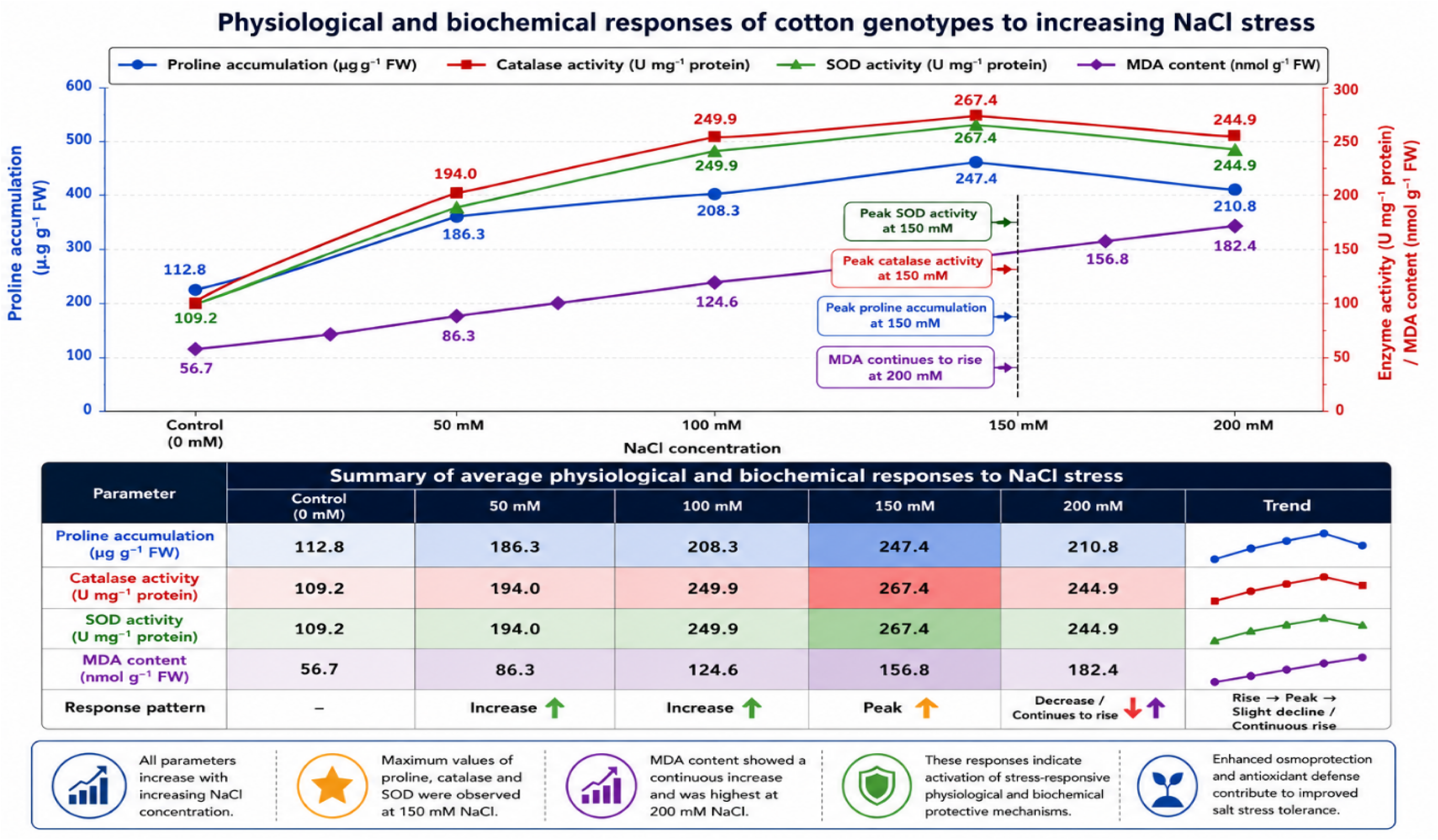
Integrated physiological and biochemical responses of cotton genotypes under increasing NaCl-induced salt stress.

Figure 6. Integrated physiological and biochemical responses of cotton genotypes exposed to increasing NaCl concentrations (0–200 mM). The figure summarizes average changes in proline accumulation, catalase (CAT) activity, superoxide dismutase (SOD) activity, and malondialdehyde (MDA) content under salt stress conditions. Progressive increases in proline accumulation and antioxidant enzyme activities were observed with increasing salinity, reaching peak values at 150 mM NaCl. In contrast, MDA content continued to increase at higher salinity levels, reflecting enhanced oxidative membrane damage under severe stress conditions. Overall, the observed patterns demonstrate the coordinated activation of protective physiological and biochemical responses in cotton under salinity stress.

Figure 7. Statistical analysis of physiological and biochemical responses of cotton genotypes exposed to different NaCl concentrations (0–200 mM). The figure presents mean values ± standard error (SE) for proline accumulation, catalase (CAT) activity, superoxide dismutase (SOD) activity, and malondialdehyde (MDA) content under salt stress conditions. Different letters above the bars/points indicate statistically significant differences between treatments according to one-way ANOVA followed by Tukey’s multiple comparison test at p < 0.05. Increasing salinity levels resulted in enhanced osmoprotective and antioxidant responses, with the highest values generally observed at 150 mM NaCl. MDA content remained elevated under severe salinity stress conditions.

**Figure 7.**
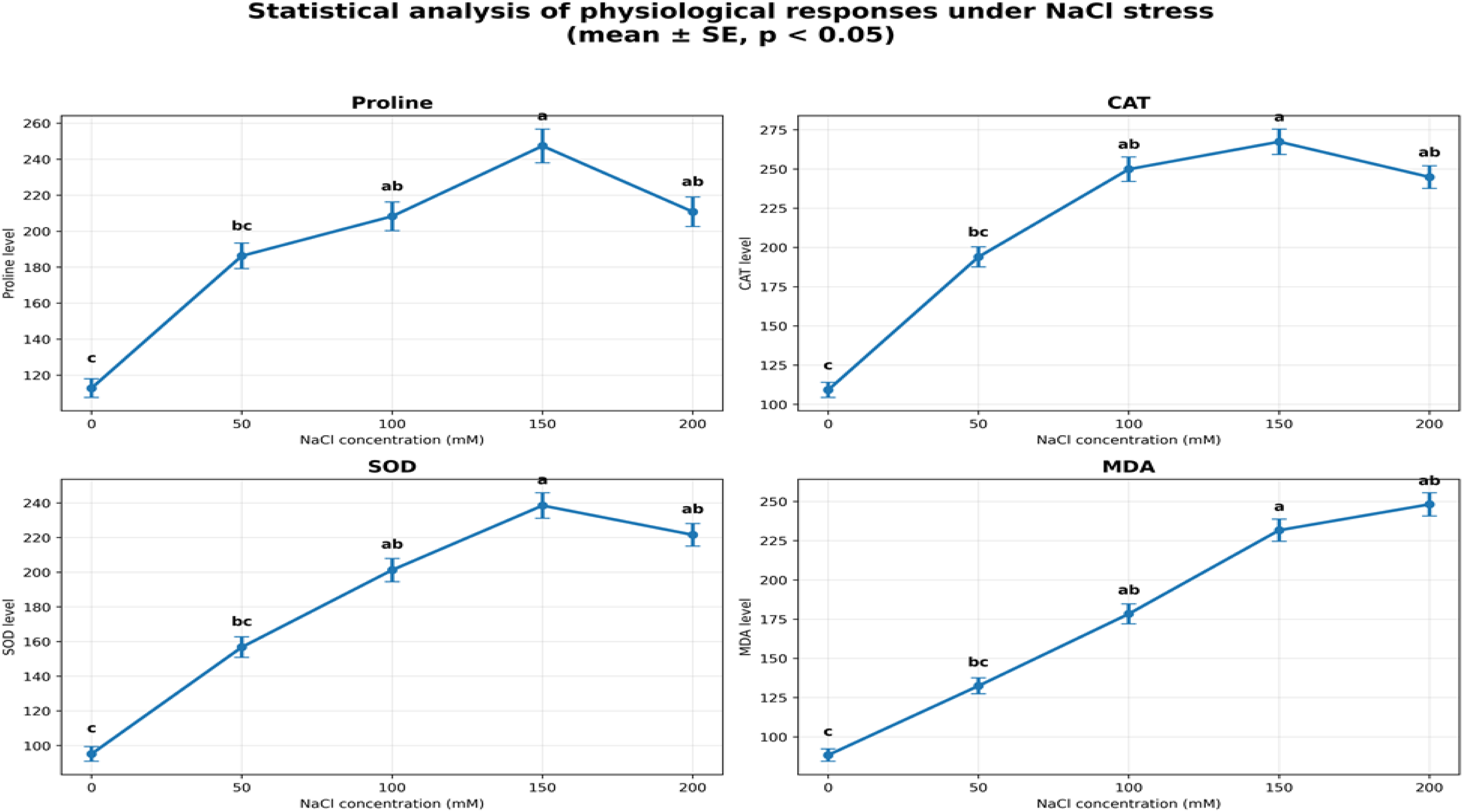
Statistical analysis of physiological and biochemical responses of cotton genotypes under NaCl-induced salt stress.

To summarize the overall physiological and biochemical responses of cotton genotypes under different salinity levels, a generalized interpretation of stress-induced changes is presented in Table 2. The table highlights the characteristic metabolic and antioxidant responses observed under control, moderate, and severe salt stress conditions.

**Table 2.** General physiological responses of cotton genotypes under different salinity levels.

| Salinity level | Physiological response |
| --- | --- |
| Control | Normal metabolic activity and stable antioxidant balance |
| Moderate salinity (100–150 mM NaCl) | Enhanced antioxidant activity and osmotic adjustment |
| Severe salinity (200 mM NaCl) | Increased oxidative stress, elevated MDA accumulation, activation of defense mechanisms |

**Values are presented as mean ± SD of three biological replicates.**

General physiological response patterns of cotton genotypes under different salinity levels are summarized in Table 2. Increasing salinity levels progressively stimulated osmotic regulation and antioxidant defense responses, while simultaneously enhancing oxidative stress development.

## DISCUSSION

Salinity stress is considered one of the most severe abiotic constraints limiting cotton productivity worldwide. Excessive accumulation of soluble salts in the rhizosphere disrupts cellular homeostasis through osmotic stress, ion toxicity, nutrient imbalance, and overproduction of reactive oxygen species (ROS), ultimately leading to oxidative damage and metabolic dysfunction. Consequently, plant tolerance to salinity largely depends on the efficiency of osmotic adjustment, antioxidant defense systems, and maintenance of membrane stability under stress conditions.

The present study demonstrated that NaCl-induced salinity stress triggered pronounced physiological and biochemical alterations in cotton genotypes, including significant changes in proline accumulation, antioxidant enzyme activities (SOD and CAT), and MDA content. However, the magnitude and pattern of these responses differed considerably among genotypes, indicating substantial genotype-dependent variability in salinity tolerance mechanisms.

One of the most prominent adaptive responses observed under saline conditions was enhanced proline accumulation. Proline is widely recognized as a multifunctional osmoprotectant involved in osmotic adjustment, stabilization of proteins and membranes, scavenging of reactive oxygen species, regulation of cellular redox balance, and protection of photosynthetic machinery during stress exposure. In the present study, the genotypes “Nasaf”, “Gulbahor-2”, “Ravnaq-1”, “Buxoro-6”, “Afsona”, “Baraka”, “Namangan-77”, “Porloq-1”, and “C-4727” exhibited comparatively greater proline accumulation, particularly under moderate and severe salinity levels. These findings suggest that enhanced osmotic adjustment capacity may contribute to improved stress adaptation in these genotypes.

The obtained results are consistent with previous reports demonstrating the protective role of proline under salinity stress. Meloni et al. (2001) observed substantial increases in proline accumulation in salt-stressed cotton plants, whereas Ashraf (2009) reported that enhanced proline biosynthesis is strongly associated with stress tolerance mechanisms in higher plants. Hosseinifard et al. (2022) further emphasized that proline contributes to membrane stabilization and protection of cellular macromolecules under abiotic stress conditions. Similarly, Zulfiqar and Ashraf (2023) highlighted the involvement of proline in reducing oxidative injury through modulation of ROS detoxification pathways. More recently, Renzetti et al. (2024) demonstrated that genes associated with proline metabolism are transcriptionally regulated during salinity stress, indicating an important role of proline-mediated signaling in plant stress adaptation.

Besides osmotic regulation, antioxidant defense mechanisms represent another critical component of plant tolerance to salinity stress. Excessive ROS generation under saline conditions can induce oxidative damage to membrane lipids, proteins, chloroplast structures, and nucleic acids. Therefore, activation of antioxidant enzymes is essential for maintaining cellular redox homeostasis and minimizing oxidative injury.

The present findings revealed that increasing NaCl concentrations generally stimulated CAT and SOD activities in most investigated cotton genotypes. However, tolerant genotypes demonstrated substantially stronger antioxidant responses compared with sensitive genotypes. In particular, “Nasaf”, “Gulbahor-2”, “Ravnaq-1”, “Buxoro-6”, “Afsona”, “Baraka”, “Namangan-77”, “Porloq-1”, and “C-4727” showed comparatively higher CAT and SOD activities under salinity stress conditions. Elevated SOD activity may indicate enhanced dismutation of superoxide radicals into hydrogen peroxide, whereas increased CAT activity reflects efficient detoxification of hydrogen peroxide into water and oxygen. Coordinated functioning of these enzymes is therefore essential for limiting oxidative stress intensity in plant tissues exposed to salinity.

The observed enhancement of antioxidant enzyme activities in tolerant genotypes is in agreement with the studies of Foyer and Noctor (2005), who described antioxidant regulation as a central component of cellular redox homeostasis under stress conditions. Tripathy and Oelmüller (2012) also reported that antioxidant enzymes play a major role in ROS detoxification during abiotic stress exposure. Gill and Tuteja (2020) emphasized that efficient antioxidant systems are closely associated with improved stress tolerance and cellular protection in plants. Similarly, Hasanuzzaman et al. (2021) demonstrated that increased CAT and SOD activities contribute to reduced oxidative membrane injury under salinity stress. Sekmen et al. (2014) further reported that salt-tolerant cotton cultivars generally exhibit stronger antioxidant enzyme activities compared with salt-sensitive cultivars.

An important aspect of the present study was the evaluation of MDA accumulation as an indicator of lipid peroxidation and oxidative membrane damage. Increased MDA content is commonly associated with excessive ROS accumulation and deterioration of membrane integrity under stress conditions. In the current investigation, NaCl stress generally caused progressive increases in MDA accumulation across most cotton genotypes. Nevertheless, the intensity of lipid peroxidation varied substantially among genotypes.

Salt-sensitive genotypes exhibited comparatively higher MDA accumulation under elevated NaCl concentrations, indicating greater oxidative membrane injury and weaker antioxidant protection capacity. In contrast, the genotypes “Porloq-1”, “Afsona”, “Buxoro-14”, and partially “Omad” maintained relatively lower MDA levels even under severe salinity stress conditions. Reduced MDA accumulation in these genotypes may reflect more efficient ROS scavenging systems, improved membrane stability, and stronger adaptive capacity under saline environments. These observations are supported by previous studies demonstrating the relationship between oxidative stress and lipid peroxidation in plants exposed to salinity. Foyer and Noctor (2005) reported that excessive ROS accumulation promotes membrane lipid peroxidation and cellular damage. Czégény et al. (2014) further demonstrated that hydrogen peroxide accumulation contributes to oxidative burst processes and membrane deterioration in stressed plants. Hasanuzzaman et al. (2021) emphasized that oxidative membrane injury under saline conditions is strongly associated with imbalance between ROS production and antioxidant defense capacity. Wang et al. (2024) also reported that efficient antioxidant systems contribute to stabilization of cellular membranes and reduction of lipid peroxidation during salinity stress, whereas Wu et al. (2015) noted that enhanced antioxidant activity is frequently associated with lower MDA accumulation under abiotic stress conditions.

The comparative behavior of tolerant and sensitive genotypes observed in the present study suggests that salinity tolerance in cotton is not determined by a single physiological parameter, but rather by coordinated interaction between osmotic adjustment mechanisms and antioxidant defense systems. Genotypes characterized by enhanced proline accumulation together with elevated CAT and SOD activities generally exhibited lower MDA accumulation, indicating more efficient protection against oxidative membrane damage. Conversely, genotypes with weaker antioxidant responses tended to accumulate higher MDA levels, reflecting greater susceptibility to oxidative stress.

The application of heatmap visualization approaches provided an effective and integrative representation of genotype-specific stress response patterns under different salinity levels. This integrated physiological and biochemical responses analysis improved the interpretation of complex biochemical interactions and enabled clearer identification of tolerant and sensitive genotypes based on coordinated physiological and antioxidant responses.

Overall, the present findings indicate that salinity tolerance in cotton is closely associated with integrated regulation of osmoprotective and antioxidant defense pathways. Enhanced proline accumulation, elevated CAT and SOD activities, and reduced MDA accumulation may therefore serve as reliable physiological and biochemical markers for screening salinity-tolerant cotton genotypes. The tolerant genotypes identified in this study may represent valuable genetic resources for future breeding programs aimed at developing cotton cultivars with improved adaptation to saline environments under increasing global environmental stress conditions.

To provide a clearer physiological interpretation of the investigated biochemical markers under salinity stress, the functional significance of each parameter is summarized in Table 3. The table highlights the relationship between antioxidant responses, osmotic regulation, and oxidative damage observed under saline conditions.

**Table 3.** Physiological interpretation of investigated biochemical markers under salinity stress.

| Biochemical parameter | Physiological role under salinity stress | Interpretation of increased values |
| --- | --- | --- |
| Proline | Osmotic adjustment and membrane stabilization | Indicates activation of osmoprotective mechanisms |
| SOD (Superoxide dismutase) | Detoxification of superoxide radicals | Reflects activation of antioxidant defense system |
| CAT (Catalase) | Decomposition of hydrogen peroxide | Indicates regulation of oxidative stress |
| MDA (Malondialdehyde) | Marker of lipid peroxidation | Reflects membrane oxidative damage intensity |

**Values are presented as mean ± standard deviation (SD) of three biological replicates.**

The physiological significance and stress-related interpretation of the investigated biochemical markers are presented in Table 3. The analyzed parameters collectively reflect osmotic regulation, antioxidant defense activity, and oxidative membrane damage under salinity stress conditions.

## CONCLUSION

The results of the present study confirmed that NaCl-induced salinity stress exerts a significant influence on the physiological and biochemical processes of cotton genotypes. Increasing NaCl concentrations were accompanied by changes in the activity of antioxidant enzymes, including superoxide dismutase (SOD) and catalase (CAT), enhanced proline accumulation, and increased malondialdehyde (MDA) content, indicating the activation of antioxidant and osmotic defense mechanisms under saline conditions.

The obtained findings revealed pronounced genotype-dependent variability in physiological and biochemical responses to salt stress. The genotypes “Nasaf”, “Gulbahor-2”, “Ravnaq-1”, “Buxoro-6”, “Afsona”, “Baraka”, “Namangan-77”, “Porloq-1”, and “C-4727” were characterized by elevated antioxidant enzyme activity, enhanced proline accumulation, and comparatively lower MDA content. These characteristics indicate more efficient antioxidant defense mechanisms, improved membrane stability, and greater adaptive capacity to salinity stress conditions.

In contrast, the genotypes “Xin Lu Zhoo-78”, “Chuntay-2”, and partially “Omad” exhibited relatively higher MDA accumulation accompanied by lower antioxidant enzyme activity, suggesting greater sensitivity to NaCl-induced oxidative stress.

The study demonstrated that proline accumulation, increased SOD and CAT activities, and reduced lipid peroxidation intensity represent important adaptive mechanisms associated with salinity tolerance in cotton. Conversely, elevated MDA content reflected the degree of oxidative membrane damage and may therefore serve as an indicator of stress sensitivity in plants.

The application of heatmap visualization approaches enabled effective characterization of genotype-specific physiological and biochemical responses under different salinity levels and significantly improved the interpretation of complex stress-response patterns.

Overall, the obtained results confirm that antioxidant enzyme activities (SOD and CAT), proline accumulation, and MDA content may be considered reliable physiological and biochemical markers for evaluating salinity tolerance in cotton genotypes. The salt-tolerant genotypes identified in this study represent valuable genetic resources for future breeding programs aimed at developing highly salinity-tolerant cotton cultivars suitable for cultivation under saline soil conditions.

## Acknowledgements

The authors gratefully acknowledge the financial support of the Academy of Sciences of the Republic of Uzbekistan and the Center for Genomics and Bioinformatics. Special thanks are extended to the Academy of Sciences for their continued support.

## Ethics Approval

Not applicable to this paper.

## Author Contributions

NRR, ASI, IBS and VSK carried out the experiments and wrote andrevised the manuscript, performed statistical analysis. SBK, FRS, JKN, MZ, ZZY and RAJ participated in the experiments, collected the data, and prepared the manuscript. ZTB edited and approved the manuscript. All authors read and approved the final manuscript.

## Conflict of interest

The authors declare that they have no competing interests.

## Data Availability

Data presented in this study will be available on a fair request to the corresponding author.

## Notes

### Competing Interest Statement

The authors have declared no competing interest.

## References

1. Hasanuzzaman, M., Bhuyan, M. H. M. B., Zulfiqar, F., Raza, A., Mohsin, S. M., Mahmud, J. A., Fujita, M., & Fotopoulos, V. (2021). Reactive oxygen species and antioxidant defense in plants under abiotic stress: Revisiting the crucial role of a universal defense regulator. Antioxidants, 10(1), 156. 10.3390/antiox10010156

2. Wang, Y., Li, X., Zhang, H., Chen, F., & Liu, J. (2024). Antioxidant defense mechanisms and membrane stability under salinity stress in crop plants. Plants, 13(2), 214. 10.3390/plants13020214

3. Maryum Z, Luqman T, Nadeem SN, Khan SMUD, Wang B, Ditta A, et al. An overview of salinity stress, mechanism of salinity tolerance and strategies for its management in cotton. Frontiers in Plant Science. 2022;13:907937. 10.3389/fpls.2022.907937

4. Ullah I, Nadeem M, Nabi HG, Shahzad N, Abbas Z, Shakoor I, et al. Molecular networks and signaling pathways governing abiotic stress tolerance in cotton: advances and perspectives. Functional & Integrative Genomics. 2026;26(1):23. 10.1007/s10142-025-01795-8

5. Foyer, C. H., & Noctor, G. (2005). Oxidant and antioxidant signalling in plants: A re-evaluation of the concept of oxidative stress in a physiological context. Plant, Cell & Environment, 28(8), 1056–1071. 10.1111/j.1365-3040.2005.01327.x

6. Tripathy, B. C., & Oelmüller, R. (2012). Reactive oxygen species generation and signaling in plants. Plant Signaling & Behavior, 7(12), 1621–1633. 10.4161/psb.22455

7. Gill SS, Tuteja N. Reactive oxygen species and antioxidant machinery in abiotic stress tolerance in crop plants. Plant Physiology and Biochemistry. 2010;48(12):909–930. 10.1016/j.plaphy.2010.08.016

8. Hasanuzzaman M, Bhuyan MHMB, Zulfiqar F, Raza A, Mohsin SM, Mahmud JA, et al. Reactive oxygen species and antioxidant defense in plants under abiotic stress: revisiting the crucial role of a universal defense regulator. Antioxidants. 2020;9(8):681. 10.3390/antiox9080681.

9. Noreen Z, Ashraf M. Assessment of variation in antioxidative defense system in salt-treated pea cultivars and its putative use as salinity tolerance markers. Journal of Plant Physiology. 2009;166(16):1764–1774. 10.1016/j.jplph.2009.05.005

10. Ashraf, M. (2009). Biotechnological approach of improving plant salt tolerance using antioxidants as markers. Biotechnology Advances, 27(1), 84–93. 10.1016/j.biotechadv.2008.09.003

11. Hosseinifard, M., Stefaniak, S., Ghorbani Javid, M., Soltani, E., Wojtyla, Ł., & Garnczarska, M. (2022). Contribution of exogenous proline to abiotic stresses tolerance in plants: A review. International Journal of Molecular Sciences, 23(9), 5186. 10.3390/ijms23095186

12. Meloni, D. A., Oliva, M. A., Martinez, C. A., & Cambraia, J. (2001). Photosynthesis and activity of antioxidant enzymes in cotton under salt stress. Biologia Plantarum, 44(1), 115–121. 10.1023/A:1017986807916

13. Zulfiqar, F., & Ashraf, M. (2023). Proline-induced abiotic stress tolerance in plants: Physiological and molecular perspectives. Journal of Plant Growth Regulation, 42(5), 2568–2584. 10.1007/s00344-022-10646-3

14. Czégény, G., Pusztahelyi, T., Hideg, É., & Kőszegi, T. (2014). Hydrogen peroxide contributes to the oxidative burst and lipid peroxidation under abiotic stress conditions in plants. Plant Physiology and Biochemistry, 80, 126–132. 10.1016/j.plaphy.2014.03.030

15. Sekmen, A. H., Türkan, İ., & Takio, S. (2014). Differential responses of antioxidative enzymes and lipid peroxidation to salt stress in salt-tolerant and salt-sensitive cotton cultivars. Plant Growth Regulation, 72(2), 123–134. 10.1007/s10725-013-9845-3

16. Ahmad P, Jaleel CA, Salem MA, Nabi G, Sharma S. Roles of enzymatic and non-enzymatic antioxidants in plants during abiotic stress. Critical Reviews in Biotechnology. 2010;30(3):161–175. doi: 10.3109/07388550903524243

17. Babadjanova FI, Ubaydullaeva KA, Rakhmanov BK, Bolkiev AA, Abdullaev AN, Abdullaev SA, Yusupov HN, Rakhmatova NR, Kushakov SO, Ayubov MS, Buriev ZT. Modern biotechnological approaches to enhance plant responses to abiotic stresses. Plant Science Today. 2026;13(1):1–9. 10.14719/pst.11797

18. Rakhmatova NR, Imamkhodjaeva AS, Kamburova VS, Salakhutdinov IB, Radjapov FS, Mamatkulova ShH, Isomiddinova OL, Usmanov DE, Norbekov JK, Ubaydullaeva KhA, Abdullaev AN, Kushakov ShO, Kadirova ShB, Khusenov NN, Kholmuradova MM, Babadjanova FI, Rakhmanov BK, Buriev ZT. Identification of the responsiveness of some enzymes of the antioxidant system of the biotechnological cotton variety to salt stress. Plant Science Today. 2025; 12(4): 1–6. 10.14719/pst.9921

19. Heath RL, Packer L. Photoperoxidation in isolated chloroplasts. I. Kinetics and stoichiometry of fatty acid peroxidation. Archives of Biochemistry and Biophysics. 1968;125(1):189–198. 10.1016/0003-9861(68)90654-1

20. Leonowicz G, Trzebuniak KF, Zimak-Piekarczyk P, Ślesak I, Mysliwa-Kurdziel B. The activity of superoxide dismutases (SODs) at the early stages of wheat deetiolation. PLoS ONE. 2018;13(3):e0194678. 10.1371/journal.pone.0194678

21. Beers RF Jr, Sizer IW. A spectrophotometric method for measuring the breakdown of hydrogen peroxide by catalase. Journal of Biological Chemistry. 1952;195(1):133–140. 10.1016/S0021-9258(19)50881-X

22. Meloni DA, Oliva MA, Martinez CA, Cambraia J. Photosynthesis and proline metabolism in cotton under salt stress. Biologia Plantarum. 2001;44(3):411–414. 10.1023/A:1012408407961

23. Renzetti, A., Garcia-Mina, J. M., & Arnao, M. B. (2024). Proline metabolism and signaling pathways under salinity stress in plants. Plant Stress, 11, 100245. 10.1016/j.stress.2023.100245

24. Sekmen AH, Türkan İ, Takio S. Differential responses of antioxidative enzymes and lipid peroxidation to salinity stress in salt-tolerant and salt-sensitive cotton cultivars. Environmental and Experimental Botany. 2014;105:32–42. 10.1016/j.envexpbot.2014.04.006

25. Wu, G. Q., Liang, N., Feng, R. J., & Zhang, J. J. (2015). Mitigation of salinity-induced oxidative damage by enhanced antioxidant enzyme activities in plants. Acta Physiologiae Plantarum, 37(4), 69. 10.1007/s11738-015-1827-5

